# Clustering of dispersal corridors in metapopulations leads to higher rates of recovery following subpopulation extinction

**DOI:** 10.1101/2020.01.29.925529

**Authors:** Helen M. Kurkjian

**Author notes:** Department of Biology, Boston College, Chestnut Hill, MA, 02467, USA.

## Abstract

Understanding how spatially divided populations are affected by the physical characteristics of the landscapes they occupy is critical to their conservation. While some metapopulations have dispersal corridors spread relatively evenly through space in a homogeneous arrangement such that most subpopulations are connected to a few neighbors, others may have corridors clustered in a heterogeneous arrangement, creating a few highly connected subpopulations and leaving most subpopulations with only one or two neighbors. Graph theory and empirical data from other biological and non-biological networks suggest that heterogeneous metapopulations should be the most robust to subpopulation extinction. Here, I used *Pseudomonas syringae* pv. *syringae* B728a in metapopulation microcosms to compare the recovery of metapopulations with homogeneous and heterogeneous corridor arrangements following small, medium, and large subpopulation extinction events. I found that while metapopulations with heterogeneous corridor arrangements had the fastest rates of recovery following extinction events of all sizes and had the shortest absolute time to recovery following medium-sized extinction events, metapopulations with homogeneous corridor arrangements had the shortest time to recovery following the smallest extinction events.

## Introduction

A metapopulation is a collection of subpopulations occupying spatially divided habitat fragments, but connected via dispersal across less adequate habitat (Hanski 1991). Understanding how patchy habitats and fragmentation affect survival, dispersal, and growth within metapopulations is critical to their conservation. Theoretical models make many assumptions about growth and dispersal to predict metapopulation dynamics (Amarasekare 1998, Parker 1999, Hanski and Ovaskainen 2000). To effectively use such models to prioritize conservation efforts, we must determine which are best supported by data using experimental tests of their predictions (Kareiva 1989, Holyoak and Lawler 2005).

Metapopulation dynamics are affected by factors beyond the within-patch survival and growth of constituent subpopulations. A growing body of literature links classic metapopulation theory with graph, or network, theory. In the latter framework, the metapopulation’s subpopulations occupy habitat fragments, which are nodes in a graph, and dispersal between any two fragments can be represented by a graph edge (Urban et al. 2009). Using this network-theoretic approach, we can use graph analytic metrics such as degree distributions and spanning trees to better describe metapopulations. One such metric is network heterogeneity, which can range from entirely homogeneous, in which all subpopulations are equally connected to their neighbors, to strongly heterogeneous, as, for example, when most subpopulations are connected to only one or two neighbors while a few “hub” subpopulations are more highly connected (Dale and Fortin 2010, Gilarranz and Bascompte 2012). In other words, the connectivity degree of each subpopulation, or the number of dispersal corridors by which it is connected to the network is uniform in a homogeneous metapopulation, but right-skewed in a more heterogeneous metapopulation (Urban and Keitt 2001, Molofsky and Ferdy 2005, Artzy-Randrup and Stone 2010).

Interpreting metapopulation dynamics through the lens of such network-theoretic metrics allows us to draw parallels among diverse types of networks, both biological and non-biological. For example, empirical work in systems as diverse as the World Wide Web (Albert et al. 2000) and stream systems (Fagan 2002) supports models suggesting that heterogeneous networks are more robust to local failure than their more homogeneous counterparts.

Theoretical modelling by Albert and colleagues (2000) demonstrated that heterogeneous networks are more robust to failures and that such robustness cannot be explained simply by path redundancy. Rather, heterogeneous networks have a smaller diameter, or average path length between any two nodes, and this interconnectedness allows information, or in a metapopulation, dispersing individuals, to cross the network more quickly. And a heterogeneous network has so few nodes that are highly connected that a random local failure is more likely to happen at a poorly connected node. Thus, the remaining nodes in such a network can still communicate with each other, which makes the network less likely to experience global failure than would a more homogeneous network experiencing a similar level of local failure.

In metapopulations, a subpopulation can be rescued from extinction by migration from neighboring populations, and such rescues can increase the probability that the metapopulation will persist through time (Brown and Kodric-Brown 1977). Subpopulation connectedness can influence how important this rescue effect is to metapopulation persistence. Because highly connected subpopulations are more likely to be rescued by migration from neighbors, a metapopulation containing many highly connected subpopulations is expected to have a greater rate of recovery and, therefore, a higher probability of persistence than a metapopulation with fewer highly connected subpopulations (Eriksson et al. 2014). However, whether the arrangement of dispersal corridors in space can itself affect the recovery of the metapopulation has rarely been explored empirically, despite the fact that a deeper understanding of such dynamics could provide critical conservation information.

In addition, the size of the extinction, or percentage of subpopulations that go extinct, could interact with the effects of corridor arrangement on metapopulation recovery. For example, Albert and colleagues found that as the fraction of nodes lost in a network rose, the diameter of the network rose slowly in more homogeneous networks. However, in heterogeneous networks, the diameter of the network was largely unaffected when nodes were lost at random, but when highly connected nodes were targeted for removal, the diameter of the network increased rapidly as the fraction of nodes lost increased.

Here, I used the network-theoretic concept of heterogeneity to characterize dispersal corridor spatial distributions of experimental bacterial metapopulations and tested the effect of that heterogeneity on recovery following subpopulation extinctions. I predicted that metapopulations with heterogeneous corridor arrangements, compared to those with homogeneous arrangements, would recover faster from subpopulation extinction events of all sizes. However, at higher extinction sizes, I predicted the difference in recovery rate between homogeneous and heterogeneous metapopulations would be larger.

## Methods

To compare the rate of recolonization and recovery following subpopulation extinction between metapopulations with different degrees of network heterogeneity in their dispersal corridors I conducted a fully crossed experiment with two treatments: corridor arrangement and extinction level. Corridor arrangement treatments were produced in Metapopulation Microcosm Plates (MMPs), which are devices that resemble 96-well microtiter plates in size and shape but with corridors connecting the wells in a desired spatial configuration. Complete methods for design, construction, and assembly of MMPs are described by Kurkjian (2019). Here, 95 of the 96 wells were connected by 176 corridors. One unconnected well served as an uninoculated control. The three corridor arrangements were: homogeneous, heterogeneous, and variable (Figure 1). In the homogeneous arrangement, wells were connected in an even lattice arrangement by 4.0 mm corridors. In the heterogeneous arrangement, wells were connected by corridors of variable lengths. Well connectivities followed a right-skewed distribution such that a few wells were highly connected whereas most were connected to only one or two other wells. In the variable arrangement, the wells were connected in the even lattice arrangement of the homogeneous treatment with the distribution of corridor lengths matching the heterogeneous treatment.

**Figure 1.**
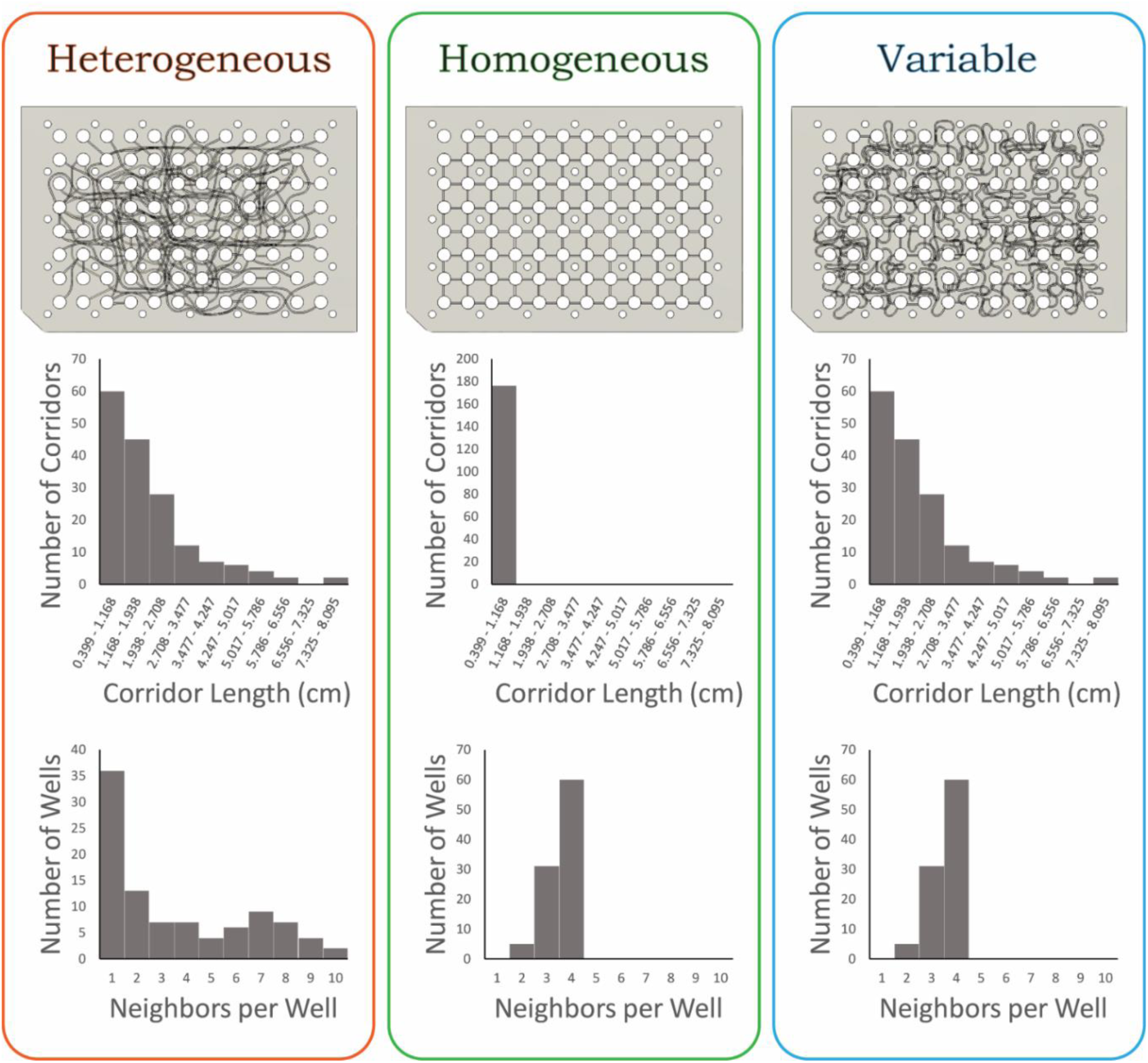
Corridor arrangement treatments: diagrams (top), histograms of corridor lengths (middle), histograms of neighbors per subpopulation (bottom).

The extinction levels were 10% (10 wells) extinct, 50% (48 wells) extinct, 90% (86 wells) extinct, and a 0% no extinction control. In each run of the experiment (Figure 2), parent plates of each corridor arrangement treatment level were needle inoculated in every well with *Pseudomonas syringae* pv. *syringae* B728a expressing green fluorescent protein from an overnight culture at a concentration of 1.0-1.7 × 10^6^ CFU/mL. The needles used for all inoculations in this experiment transferred 0.651 ± 0.347 (SD) µL volume, meaning that each well of the parent plates began with approximately 2.2-12.1 × 10^3^ CFU/mL in each well. Parent plates were incubated at 22°C for 48 hours, then subsampled into daughter plates with identical corridor arrangements using a plate replicator from which a random subset of pins had been removed (e.g. to create the 10% treatment, 10 pins were removed from the plate replicator). The identities of the extinct wells were chosen by sorting the wells of the heterogeneous corridor arrangement by their degree (number of neighbors) and choosing a random subset of each degree, approximately in proportion with the overall degree distribution of the heterogeneous treatment (Figure S1). The same well identities were extinguished in every corridor arrangement treatment, such that if well B6 was chosen as an extinct well, it was made to go extinct in all plates. Each daughter plate was incubated at 22°C for 156 hours and its fluorescence was measured in a microplate reader every 12 hours. The full experiment was run six times for a total of six replicates (plates) of each treatment combination. All uninoculated controls remained sterile.

**Figure 2.**
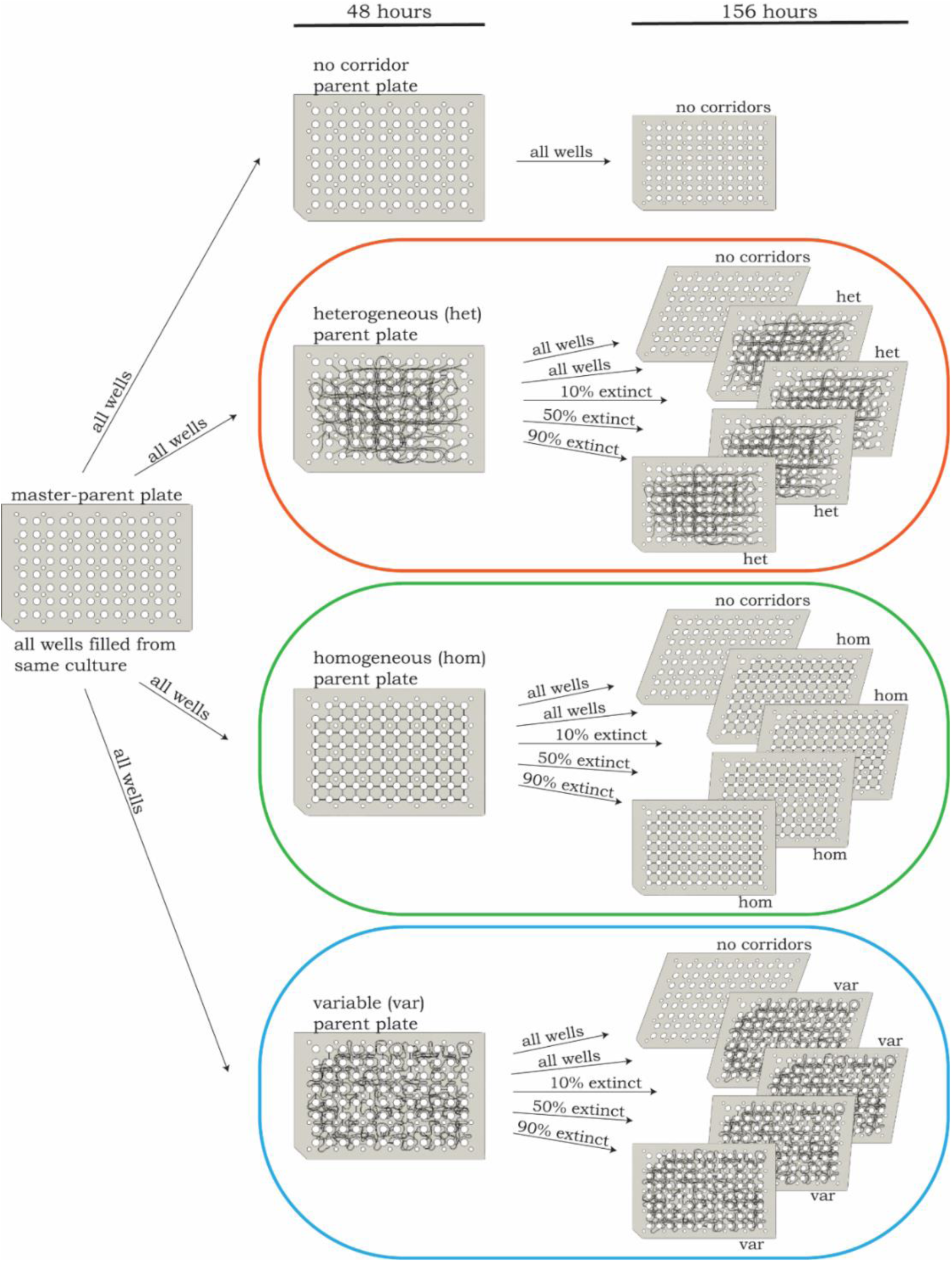
Experimental design: For each run of the experiment, all wells of the master-parent plate were filled from an overnight culture of *Pseudomonas syringae*. All wells of the master-parent plate were then sub-sampled into parent plates with homogeneous, heterogeneous, and variable corridor arrangements. These parent plates were incubated at 22°C for 48 hours. To create the different extinction levels, each parent plate was subsampled to daughter plates with identical corridor arrangements with either 10%, 50%, or 90% of plate replicator pins removed. The daughter plates were incubated 22°C for 156 hours and each was read in a plate reader every 12 hours.

To measure recovery following extinction, I calculated the deviation of each well on each treatment plate from the corresponding well on its control plate (the 0% extinction plate subsampled from the same parent plate), normalized to the control value and starting inoculum concentration, using the following formula:

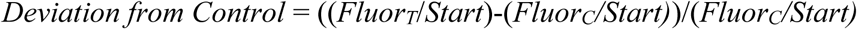

where *Fluor*_*T*_ is the fluorescence of the treatment well, *Fluor*_*C*_ is the fluorescence of the corresponding control well, and *Start* is the concentration of the culture from which the parent plate was inoculated (Figures S2 and S3). I used this quantity to calculate three recovery metrics:

1. Time to Begin Recovery Phase – I defined the recovery phase of each plate’s time series as the portion during which the change in deviation from the control was positive (i.e. the treatment plate was becoming more similar to the control). I calculated the time at which this recovery phase began for each plate.
2. Maximum Rate of Recovery – I fit each recovery phase with linear, quadratic, cubic, and quartic regressions and chose the model which fit the data best using AICc model selection. I then found the derivative of each best fit model and used it to calculate the maximum rate of recovery during the recovery phase for each treatment.
3. Time to Recovery – Using a planned contrast at each time step, I defined the time to recovery as the first time period at which there was no significant difference between the treatment and control.

I calculated all three metrics for each combination of corridor arrangement and extinction level both for the mean of all subpopulations and for extinction subpopulations only. To assess whether variability in re-colonization affected the recovery of the metapopulations, I also calculated the standard deviation of Deviation of Control of the wells of each plate.

Finally, with all subpopulations, I fit several multiple polynomial linear mixed effects models using Deviation from Control after 36 hours (to include only time points after recovery had begun for all plates) as the response variable and including run of the experiment as a random effect. Hours post-extinction, hours post-extinction squared, and hours post-extinction cubed were included as explanatory variables in all models. One or more of the following were also included in each model as explanatory variables: corridor arrangement, extinction level, and the interaction between corridor arrangement and extinction level. I chose the model with the lowest AIC_c_ value as the best fit model. I repeated this procedure with extinct subpopulations only. I assessed homogeneity of variance and normality of residuals by plotting the residuals. All analyses were performed in R (R Core Team 2016).

In all runs of this experiment, MMPs were filled with Luria-Bertani liquid medium (LB; Cold Spring Harbor Protocols 2006) containing 40 µg/mL nitrofurantoin and 15 µg/mL tetracycline to select against loss of mutant plasmids. This experiment was performed using wild type *Pseudomonas syringae* pv. *syringae* B728a containing the *pkln42gfp* plasmid, which constitutively expresses *gfp* (Dulla and Lindow 2008). This strain is motile and doubles approximately every 3 h at 15°C in this set-up. This strain was generously provided to me by the Lindow Lab, UC Berkeley.

## Results

### Time to Begin Recovery Phase

Apart from the homogeneous corridor 10% extinction treatment combination, which grew faster than the control during all periods, all 10% and 50% extinction metapopulations began to recover (approach their controls) after 24 hours. All 90% extinction metapopulations began to recover after 36 hours (Table S1a). All extinct subpopulations in the 10% and 50% extinction metapopulations began to recover after 24 hours and all extinct subpopulations in the 90% extinction metapopulations began to recover after 36 hours (Table S1b).

In general, the metapopulations experiencing 10% extinction recovered to the highest mean subpopulation sizes (relative to their controls), 50% extinction had intermediate subpopulation sizes, and the 90% extinction treatment had the lowest subpopulations sizes. Within the 10% extinction treatment, metapopulations with homogeneous corridors reached the highest subpopulations sizes, while in the 50% extinction treatment, metapopulations with heterogeneous corridors were highest (Figure 3a). The patterns were the same amongst the extinct wells (Figure 3b). Standard deviation of the “deviation from control” response variable was highest in the 90% extinction treatment, intermediate in the 50% extinction treatment, and lowest in the 10% extinction treatment. Within the 10% extinction treatment, the standard deviation was highest in the variable corridor metapopulations, intermediate in heterogeneous corridors, and lowest in homogeneous. Within the 50% and 90% extinction treatments, the standard deviation was highest in heterogeneous metapopulations, intermediate in variable metapopulations, and lowest in homogeneous (Figure 4a). The patterns were generally the same amongst extinct wells, with the exception of the heterogeneous 50% extinction treatment combination, which had the highest standard deviation (Figure 4b).

**Figure 3.**
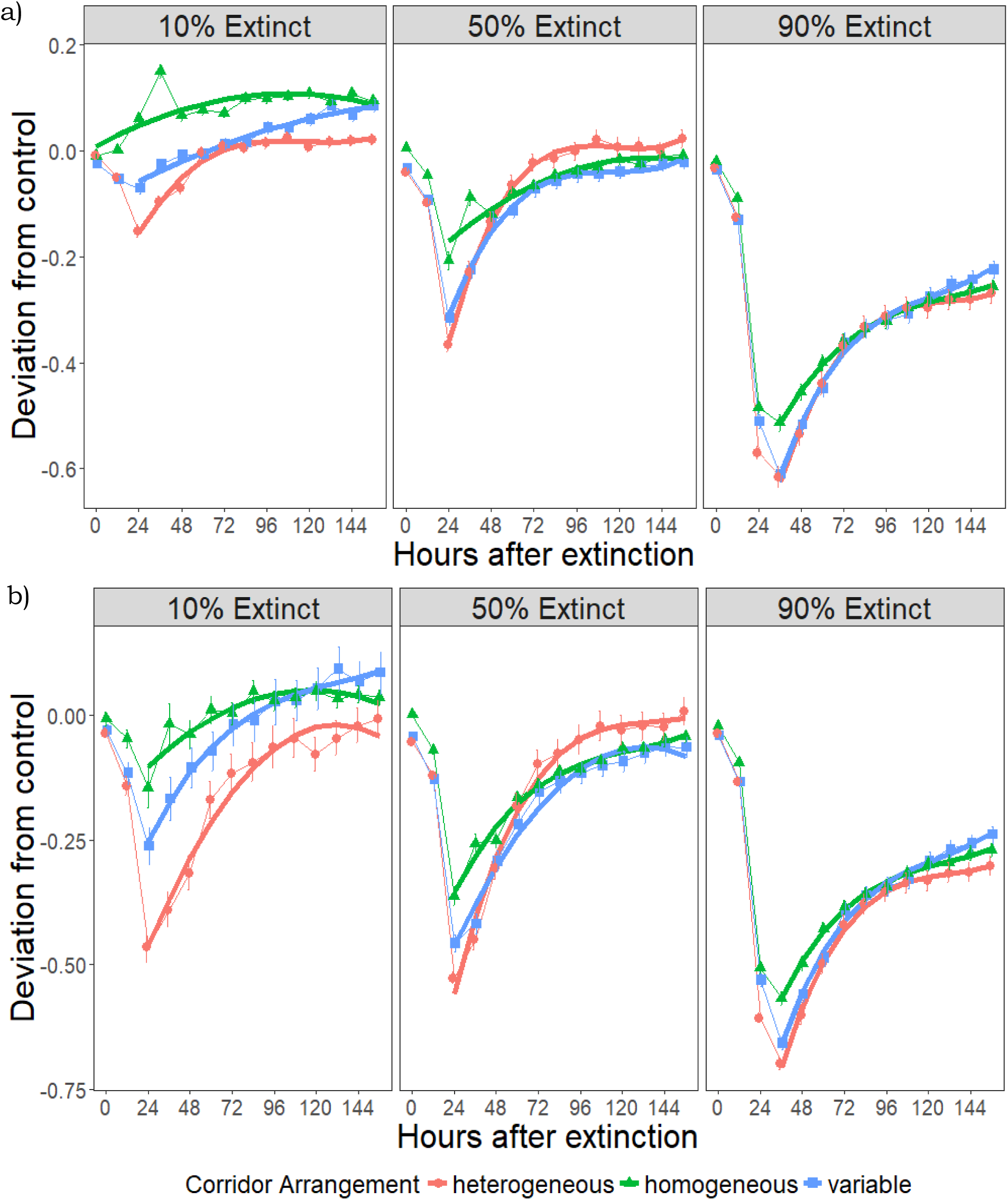
Deviation of treatment plates from control plates with heterogeneous, homogeneous, or variable corridor arrangements following 10%, 50%, or 90% extinction of a) all subpopulations or b) extinct subpopulations only. Error bars are standard error of the mean (n=6). Thick lines are best fit lines for each recovery trajectory.

**Figure 4.**
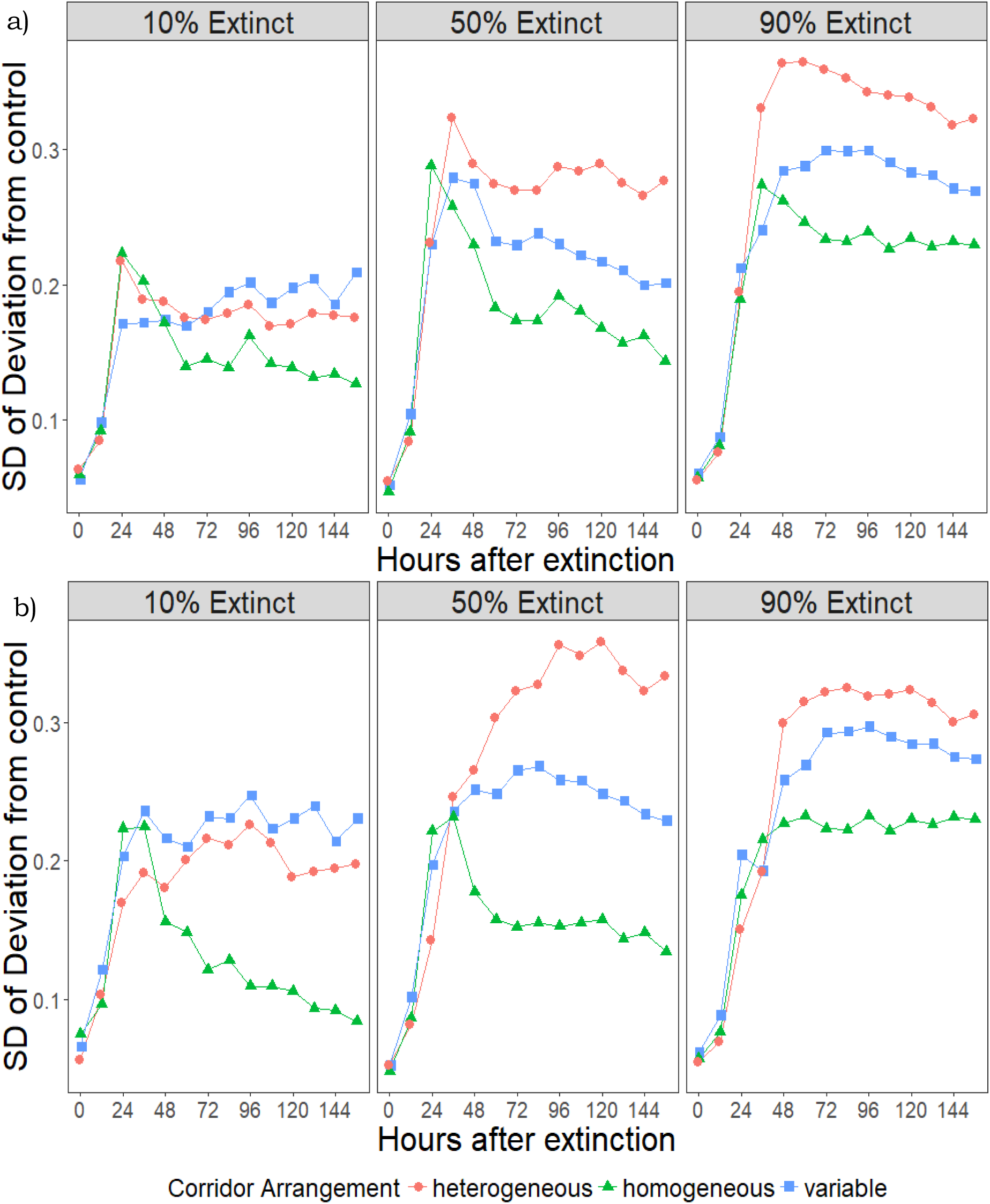
Standard deviation of “deviation from control” response variable for heterogeneous, homogeneous, or variable corridor arrangements following 10%, 50%, or 90% extinction of a) all subpopulations or b) extinct subpopulations only.

### Maximum Rate of Recovery

All recovery phase trajectories were fit best (had the lowest AIC_c_ value) by a quadratic or cubic model. The mean recoveries of all subpopulations were fit best by cubic models for all treatment combinations except 10% and 50% extinction in homogeneous corridors and 10% extinction in variable corridors, which were fit best by quadratic models (Table S2a). The recoveries of extinct subpopulations were fit best by cubic models for all treatment combinations except 10% extinction in homogeneous and heterogeneous corridors and 50% extinction in variable corridors; those exceptions were fit best by quadratic models (Table S2b).

For the mean recovery of all subpopulations, the maximum rate of recovery in metapopulations with heterogeneous corridors was highest in the 50% extinction treatment, while in metapopulations with homogeneous and variable corridors the rate was highest in the 90% extinction treatment. At all extinction levels, recovery was fastest in metapopulations with heterogeneous corridors (Figure 5a). Recovery amongst extinct subpopulations in metapopulations with heterogeneous corridors was fastest in the 50% extinction treatment. In metapopulations with homogeneous or variable corridors, recovery was fastest in the 90% extinction treatment. At all extinction levels, recovery of extinct wells was fastest in metapopulations with heterogeneous corridor arrangements (Figure 5b).

**Figure 5.**
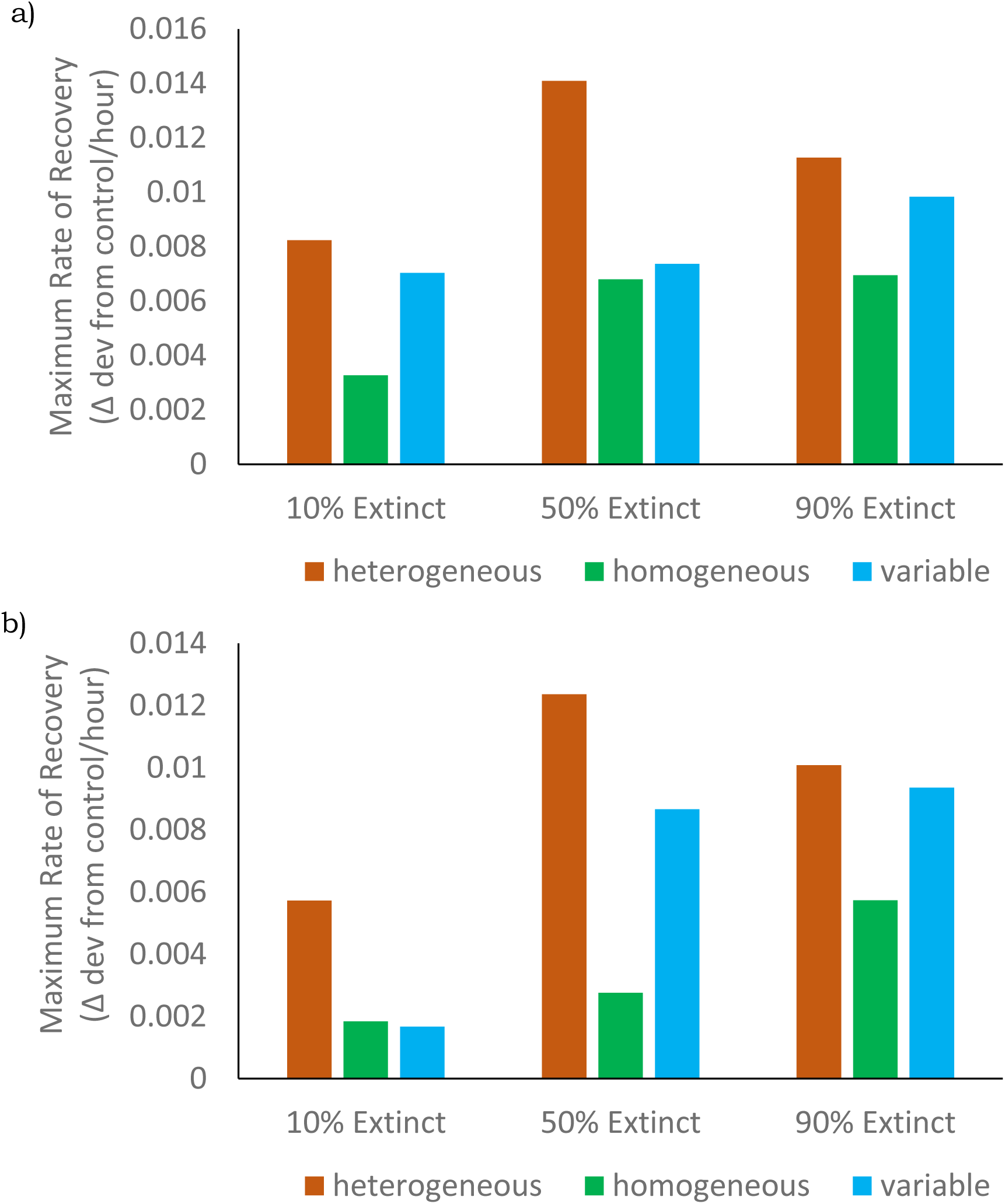
Maximum rate of recovery, measured as change in “deviation from control” response variable per hour, for each combination of corridor arrangement and extinction level for a) all subpopulations and b) extinct subpopulations only. This recovery rate maximum was calculated by finding the maximum derivative of the best-fit model [see Table S2] during the recovery phase.

### Time to Recovery

In the 10% extinction treatment, metapopulations with a heterogeneous corridor arrangement became statistically indistinguishable from their controls after 60 hours, whereas homogeneous metapopulations took only 12 hours, and metapopulations with the variable corridor arrangement took 36 hours. In the 50% extinction treatment, heterogeneous metapopulations still took 60 hours to recover to the control condition, whereas homogeneous metapopulations took 108 hours, and metapopulations with the variable corridor arrangement took 72 hours. No metapopulations recovered to the control condition following a 90% extinction (Figure 6a).

**Figure 6.**
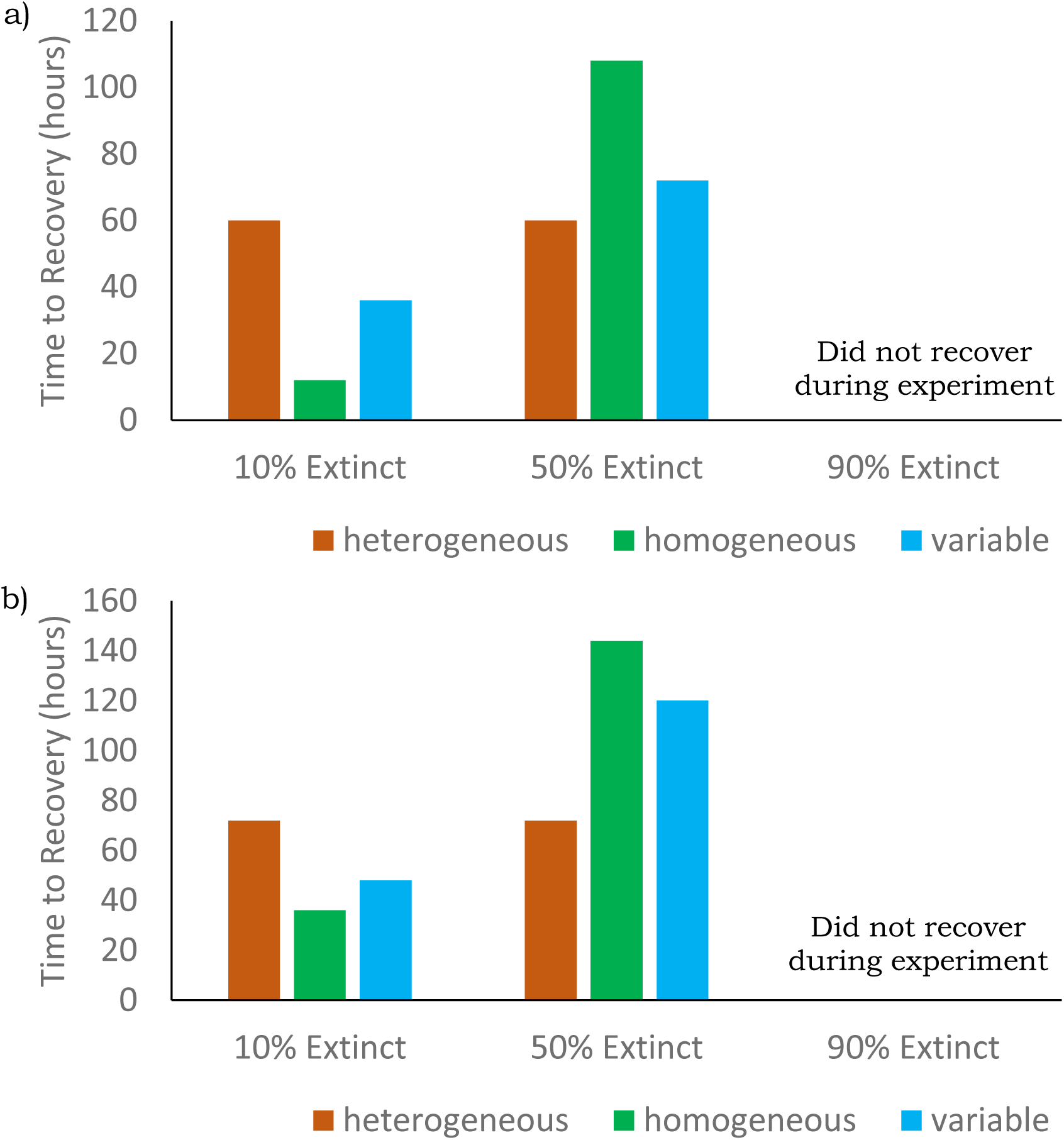
Hours to recovery for each corridor arrangement by extinction level treatment combination for a) all subpopulations and b) extinct subpopulations only.

In metapopulations with the heterogeneous corridor arrangement, extinct subpopulations took an average of 72 hours to recover from both 10% and 50% extinction treatments. In metapopulations with the homogeneous corridor arrangement, extinct subpopulations took 36 hours and 144 hours to recover from 10% and 50% extinction treatments, respectively. In metapopulations with variable corridor arrangements, extinct subpopulations took 48 hours and 120 hours to recover from 10% and 50% extinction treatments, respectively (Figure 6b).

### Polynomial Regression

All polynomial regression models of all subpopulation recoveries containing an interaction term fit the data equivalently well and better than all models that did not contain an interaction term (Table S3a). Tukey’s Honestly Significant Difference showed that all corridor arrangement by extinction level treatment combinations were significantly different from all others (p_HSD_ < 0.001) except 50% extinction in homogeneous corridors from 50% extinction in heterogeneous corridors (p_HSD_ = 0.149), 90% extinction in variable corridors from 90% extinction in homogeneous corridors (p_HSD_ = 0.102), and 90% extinction in variable corridors from 90% extinction in heterogeneous corridors (p_HSD_ = 0.173). The same pattern was true among models of extinct subpopulation recoveries (Table S3b).

## Discussion

In this experiment, metapopulations with heterogeneous corridor arrangements had a faster maximum rate of recovery from all levels of extinction than those with other corridor arrangements, which matched my predictions. However, that greater maximum speed translated to a shorter absolute time to recovery only in the 50% extinction treatment. Following the 10% extinction treatment, metapopulations with homogeneous corridor arrangements recovered to their control condition in a shorter absolute time than did the heterogeneous metapopulations. This may be because when extinctions occur in a random subset of subpopulations, few are likely to occur in adjacent subpopulations, and therefore an extinct subpopulation is almost certain to be adjacent to an occupied subpopulation, which leads to rapid recolonization. In the homogeneous treatment, that extinct subpopulation is always only a short distance from its occupied neighbor, again leading to rapid recolonization. However, in the heterogeneous treatment, bacteria may have to travel down a longer corridor, despite being only a short straight-line distance from the adjacent extinct subpopulation. Most work on network heterogeneity, including that of Albert and colleagues (2000), considers unweighted networks, in which all edges are equal. In metapopulation ecology, however, the edges of spatial networks are weighted. Assuming that the time it takes an individual organism to travel from one patch to another is a function of the length of corridor between the patches, each dispersal corridor could be assigned a weight proportional to its length (Urban et al. 2009). Therefore, predictions that heterogeneous networks will be the most robust to node failure in unweighted networks may not translate perfectly to recovery dynamics in weighted networks.

To further examine this idea, we can compare the recovery of metapopulations with the variable corridor arrangement to that of metapopulations with heterogeneous and homogeneous corridor arrangements. The variable treatment had a corridor arrangement in a regular lattice which matched that of the homogeneous treatment, but with a distribution of corridor lengths which matched that of the heterogeneous treatment. If the differences in recovery rate and time between the heterogeneous and homogeneous treatments can be explained primarily by the addition of longer dispersal corridors in the heterogeneous treatment, then recovery in the variable treatment should be similar to that of the heterogeneous treatment. If instead those differences are due primarily to differences in the corridor arrangement, then recovery in the variable treatment should be more similar to that of the homogeneous treatment. With one exception, both the rates of recovery and the absolute times to recovery of the variable treatment are intermediate between those of the heterogeneous and homogeneous treatments at every extinction level. The exception is that the recovery rate of all subpopulations in the 10% extinction treatment was slightly lower in metapopulations with a variable corridor arrangement than in those with either of the other corridor treatments. This result suggests that, while some of the difference between the heterogeneous and homogeneous treatments can be attributed to their differences in corridor length, a portion can also be attributed directly to differences in their network heterogeneity.

In standard deviation too, the variable corridor treatment is intermediate between the homogeneous corridors, which had the lowest variation in deviation from the control, and heterogeneous corridors, which had the highest variation, except for at the 10% extinction level when the variable treatment slightly exceeded the heterogeneous in variability. Here again, it is possible that the difference in corridor length between the homogeneous and heterogeneous treatments contributes to the difference in their variability, because in order for an extinct subpopulation to be recolonized individuals must travel from an occupied subpopulation. In a homogeneous metapopulation that distance is always the same, while in a heterogeneous metapopulation the distance, and therefore the time for an extinct subpopulation to be recolonized, will vary. If this variability in corridor length and travel time was the principal driver of differences in recovery between the heterogeneous and homogeneous corridor treatments, we would expect the standard deviation of the variable treatment to match that of the heterogeneous treatment. Its position intermediate between the heterogeneous and homogeneous treatments suggests again that while corridor length contributes to this difference, network heterogeneity is also important.

While it is unfortunate that none of the 90% extinction treatments recovered fully to the level of their controls within the timeframe of the experiments (which due to the physical constraints of the experimental setup had to be confined to 5 days), we can consider their rates of recovery. In the 10% extinction treatment, despite the fact that the heterogeneous corridor metapopulation has the highest rate of recovery, the homogeneous corridor metapopulation recovers in the shortest absolute time because it begins its recovery sooner and never deviates as far from its control as does the heterogeneous metapopulation. In the 50% extinction treatment, the heterogeneous metapopulation again deviates the farthest from its own control, but the combination of beginning its recovery at the same time as the other corridor arrangements and having a faster rate of recovery, leads to a shorter absolute time to recovery. In the 90% extinction treatment, by comparison, the heterogeneous metapopulation continues its pattern of deviating the farthest from its control, although the difference between the greatest deviation of the homogeneous and heterogeneous treatments is smaller than in the other extinction treatments. The heterogeneous metapopulation also has the highest rate of recovery in the 90% extinction treatment, although it is lower than the rate of recovery of the heterogeneous metapopulation in the 50% extinction treatment and the difference between the rates of the heterogeneous and homogeneous metapopulations in the 90% extinction treatment is much smaller than in the 50% extinction treatment. This suggests that while the heterogeneous metapopulation had the shortest time to recovery in the 50% extinction treatment, that pattern might not have continued in the 90% extinction treatments, had they had the time and resources to recover to control levels. This switch in order of recovery from homogeneous soonest in the 10% extinction treatment, to heterogeneous soonest in the 50% extinction treatment, and possibly back again in the 90% extinction treatment, may explain why the data was fit equally well by all polynomial models that contained a corridor arrangement by extinction level interaction term, meaning that all recovery trajectories were different from each other with no clear trends in the main effects of corridor arrangement or extinction level.

The effects of rescue by recolonization from neighboring subpopulations are important for long-term metapopulation persistence. For example, and Gilarranz and Bascompte (2012) used variations of the Hanski and Ovaskainen (2000) and Levins (1969) models to predict that higher network heterogeneity in metapopulations will lead to a higher proportion of occupied patches, but that that pattern will reverse at low extinction-to-colonization ratios. Highly connected subpopulations are likely to be recolonized more quickly by their occupied neighbors. But whether that higher rate of recolonization translates into faster recovery of the metapopulation has not been explored empirically. In this experiment, extinct subpopulations were recolonized fastest in metapopulations with heterogeneous corridors, and that recolonization led to faster recovery of the metapopulations themselves.

Understanding this type of interplay between connectivity and metapopulation dynamics is critical to conservation. For example, Fortuna and colleagues (Fortuna et al. 2006) found that variation in the wetness of the environment affected the availability of dispersal corridors for amphibians between temporary ponds, but that because of underlying connectivity amongst the ponds, amphibian dispersal was unlikely to be dramatically affected by loss of individual ponds. Similarly, Cowley, Johnson, and Pocock (2015) used network-theoretic metrics to model the spread of oak processionary moths through oak woodlands and identify the most important patches or ‘pinch points’ of invasion. Their results suggested that the patches they identified would be most critical for conservation interventions to prevent invasion spread.

A better understanding of how the spatial arrangement of dispersal corridors on the landscape might lead to different recovery outcomes depending on the size of the of the extinction event could be useful in guiding efforts to plan the positions of artificial dispersal corridors between habitat patches or preserve existing corridors. This experiment demonstrates that heterogeneous connectivity among subpopulations can lead to faster recovery following subpopulation extinction, but that greater speed of recovery may not always translate to a shorter absolute time to recovery.

## Acknowledgements

I thank Ellen Simms for her guidance in all stages of this project, members of the Simms Lab for their many helpful comments, Monica Hernandez and Tyler Helmann for bacterial cultures and advice, Caroline Williams and the Williams Lab for sharing equipment and assistance, Teffany Bareng, Monica Sadhu, and Nicholas Jourjine for their assistance in the lab, and David Ackerly, Laurel Larsen, and Steven Lindow for their helpful discussions. This work was supported by a National Science Foundation Doctoral Dissertation Improvement Grant [DEB-1601762].

## Literature Cited

Albert, R., H. Jeong, and A.-L. Barabási. 2000. Error and attack tolerance of complex networks. Nature 406:378.

Amarasekare, P. 1998. Allee Effects in Metapopulation Dynamics. The American Naturalist 152:298–302.

Artzy-Randrup, Y., and L. Stone. 2010. Connectivity, Cycles, and Persistence Thresholds in Metapopulation Networks. PLOS Computational Biology 6.

Brown, J. H., and A. Kodric-Brown. 1977. Turnover Rates in Insular Biogeography: Effect of Immigration on Extinction. Ecology 58:445–449.

Cold Spring Harbor Protocols. 2006. LB (Luria-Bertani) liquid medium. Cold Spring Harbor Protocols 2006.

Cowley, D. J., O. Johnson, and M. J. O. Pocock. 2015. Using electric network theory to model the spread of oak processionary moth, Thaumetopoea processionea, in urban woodland patches. Landscape Ecology 30:905–918.

Dale, M. R. T., and M.-J. Fortin. 2010. From Graphs to Spatial Graphs. Annual Review of Ecology, Evolution, and Systematics 41:21–38.

Dulla, G., and S. Lindow. 2008. Quorum size of Pseudomonas syringae is small and dictated by water availability on the leaf surface. Proceedings of the National Academy of Sciences 105:3082–3087.

Eriksson, A., F. Elías-Wolff, B. Mehlig, and A. Manica. 2014. The emergence of the rescue effect from explicit within- and between-patch dynamics in a metapopulation. Proceedings of the Royal Society B: Biological Sciences 281.

Fagan, W. F. 2002. Connectivity, Fragmentation, and Extinction Risk in Dendritic Metapopulations. Ecology 83:3243–3249.

Fortuna, M. A., C. Gómez-Rodríguez, and J. Bascompte. 2006. Spatial network structure and amphibian persistence in stochastic environments. Proceedings of the Royal Society B: Biological Sciences 273:1429.

Gilarranz, L. J., and J. Bascompte. 2012. Spatial network structure and metapopulation persistence. Journal of Theoretical Biology 297:11–16.

Hanski, I. 1991. Single-species metapopulation dynamics: concepts, models and observations. Biological Journal of the Linnean Society 42:17–38.

Hanski, I., and O. Ovaskainen. 2000. The metapopulation capacity of a fragmented landscape. Nature 404:755.

Holyoak, M., and S. P. Lawler. 2005. The Contribution of Laboratory Experiments on Protists to Understanding Population and Metapopulation Dynamics. Pages 245–271 Advances in Ecological Research. Academic Press.

Kareiva, P. 1989. Renewing the dialogue between theory and experiments in population ecology. Pages 68–88 Perspectives in Ecological Theory. Princeton University Press.

Kurkjian, H. M. 2019. The Metapopulation Microcosm Plate: A modified 96-well plate for use in microbial metapopulation experiments. Methods in Ecology and Evolution 10:162–168.

Levins, R. 1969. Some Demographic and Genetic Consequences of Environmental Heterogeneity for Biological Control. Bulletin of the Entomological Society of America 15:237–240.

Molofsky, J., and J.-B. Ferdy. 2005. Extinction dynamics in experimental metapopulations. Proceedings of the National Academy of Sciences of the United States of America 102:3726.

Parker, M. A. 1999. Mutualism in Metapopulations of Legumes and Rhizobia. The American Naturalist 153:S48–S60.

R Core Team. 2016. R: A Language and Environment for Statistical Computing. R Foundation for Statistical Computing, Vienna, Austria.

Urban, D., and T. Keitt. 2001. Landscape Connectivity: A Graph-Theoretic Perspective. Ecology 82:1205–1218.

Urban, D., E. Minor, E. Treml, and R. Schick. 2009. Graph models of habitat mosaics. Ecology Letters 12:260–273.

