## supplemental files for "Clustering of dispersal corridors in metapopulations leads to higher rates of recovery following subpopulation extinction"

Table S1. Hours to beginning of recovery phase for each corridor arrangement by extinction level treatment combination for a) all subpopulations and b) extinct subpopulations only (no variation between replicates).

a)

| Corridor Arrangement | Extinction Level | | |
|---|---|---|---|
| | 10% | 50% | 90% |
| Heterogeneous | 24 | 24 | 36 |
| Homogeneous | 0 | 24 | 36 |
| Variable | 24 | 24 | 36 |

b)

| Corridor Arrangement | Extinction Level | | |
|---|---|---|---|
| | 10% | 50% | 90% |
| Heterogeneous | 24 | 24 | 36 |
| Homogeneous | 24 | 24 | 36 |
| Variable | 24 | 24 | 36 |

Table S2. Akaike's Information Criterion (corrected) for quadratic, cubic, and quartic fits of deviation from control (response variable) to hours post-extinction (explanatory variable) for each corridor arrangement by extinction level treatment combination for a) all subpopulations and b) extinct subpopulations only. Asterisks indicate lowest $AIC_c$ value for each treatment combination.

a)

| Corridor Arrangement | | Model | Extinction Level 10% | 50% | 90% |
|---|---|---|---|---|---|
| Corridor Arrangement | Heterogeneous | Linear | -39.4617 | -22.0386 | -24.9291 |
| | | Quadratic | -53.7318 | -39.2041 | -44.7481 |
| | | Cubic | -59.7019 * | -65.2271 * | -56.6741 * |
| | | Quartic | -51.2729 | -58.1565 | -48.1174 |
| | Homogeneous | Linear | -48.2980 | -43.9183 | -39.4198 |
| | | Quadratic | -49.3780 * | -47.0253 * | -60.1964 |
| | | Cubic | -46.6867 | -43.4258 | -67.6496 * |
| | | Quartic | -42.4921 | -36.5383 | -58.0035 |
| | Variable | Linear | -63.9602 | -31.3703 | -29.7178 |
| | | Quadratic | -64.6027 * | -46.8633 | -42.8304 |
| | | Cubic | -58.9160 | -63.3159 * | -49.0503 * |
| | | Quartic | -54.6492 | -59.3679 | -38.5186 |

b)

| Corridor Arrangement | | Model | Extinction Level 10% | 50% | 90% |
|---|---|---|---|---|---|
| | Heterogeneous | Linear | -22.6111 | -16.9759 | -22.6932 |
| | | Quadratic | -37.4289 * | -37.9181 | -43.1829 |
| | | Cubic | -36.9965 | -39.8914 * | -58.4686 * |
| | | Quartic | -34.0553 | -39.1637 | -49.7853 |
| | Homogeneous | Linear | -37.8690 | -37.1679 | -35.4313 |
| | | Quadratic | -44.2283 * | -51.3898 | -54.1195 |
| | | Cubic | -42.1109 | -51.5477 * | -63.7529 * |
| | | Quartic | -35.3900 | -43.0327 | -55.1379 |
| | Variable | Linear | -36.2196 | -26.3050 | -28.4031 |
| | | Quadratic | -53.7085 | -45.9971 * | -42.7694 |
| | | Cubic | -53.7487 * | -45.4415 | -47.7391 * |
| | | Quartic | -49.0770 | -38.8143 | -37.0167 |

Table S3. Akaike's Information Criterion (corrected) for multiple polynomial regression fits of deviation from control (response variable), linear, squared, and cubic terms for hours post-extinction (HPE), and corridor arrangement (Corr), extinction level (Ext), and corridor arrangement by extinction level interaction (Corr:Ext) for a) all subpopulations and b) extinct subpopulations only. Asterisks indicate best-fit models.

a)

| Model (Right-hand side) | Parameters | Log likelihood | $AIC_c$ |
|---|---|---|---|
| HPE + HPE^2 + HPE^3 + Corr + Ext + Corr:Ext | 14 | 61.68 | -95.34 * |
| HPE + HPE^2 + HPE^3 + Corr + Ext | 10 | -132.78 | 285.56 |
| HPE + HPE^2 + HPE^3 + Corr | 8 | -11263.56 | 22543.12 |
| HPE + HPE^2 + HPE^3 + Ext | 8 | -269.26 | 554.53 |
| HPE + HPE^2 + HPE^3 + Corr + Corr:Ext | 14 | 61.68 | -95.34 * |
| HPE + HPE^2 + HPE^3 + Ext + Corr:Ext | 14 | 61.68 | -95.34 * |
| HPE + HPE^2 + HPE^3 + Corr:Ext | 14 | 61.68 | -95.34 * |
| HPE + HPE^2 + HPE^3 | 6 | -11355.62 | 22723.25 |

b)

| Model (Right-hand side) | Parameters | Log likelihood | AIC$_c$ |
|---|---|---|---|
| HPE + HPE^2 + HPE^3 + Corr + Ext + Corr:Ext | 14 | -2070.66 | 4169.34 * |
| HPE + HPE^2 + HPE^3 + Corr + Ext | 10 | -2158.35 | 4336.70 |
| HPE + HPE^2 + HPE^3 + Corr | 8 | -5564.44 | 11144.88 |
| HPE + HPE^2 + HPE^3 + Ext | 8 | -2200.43 | 4416.86 |
| HPE + HPE^2 + HPE^3 + Corr + Corr:Ext | 14 | -2070.66 | 4169.34 * |
| HPE + HPE^2 + HPE^3 + Ext + Corr:Ext | 14 | -2070.66 | 4169.34 * |
| HPE + HPE^2 + HPE^3 + Corr:Ext | 14 | -2070.66 | 4169.34 * |
| HPE + HPE^2 + HPE^3 | 6 | -5597.59 | 11207.18 |

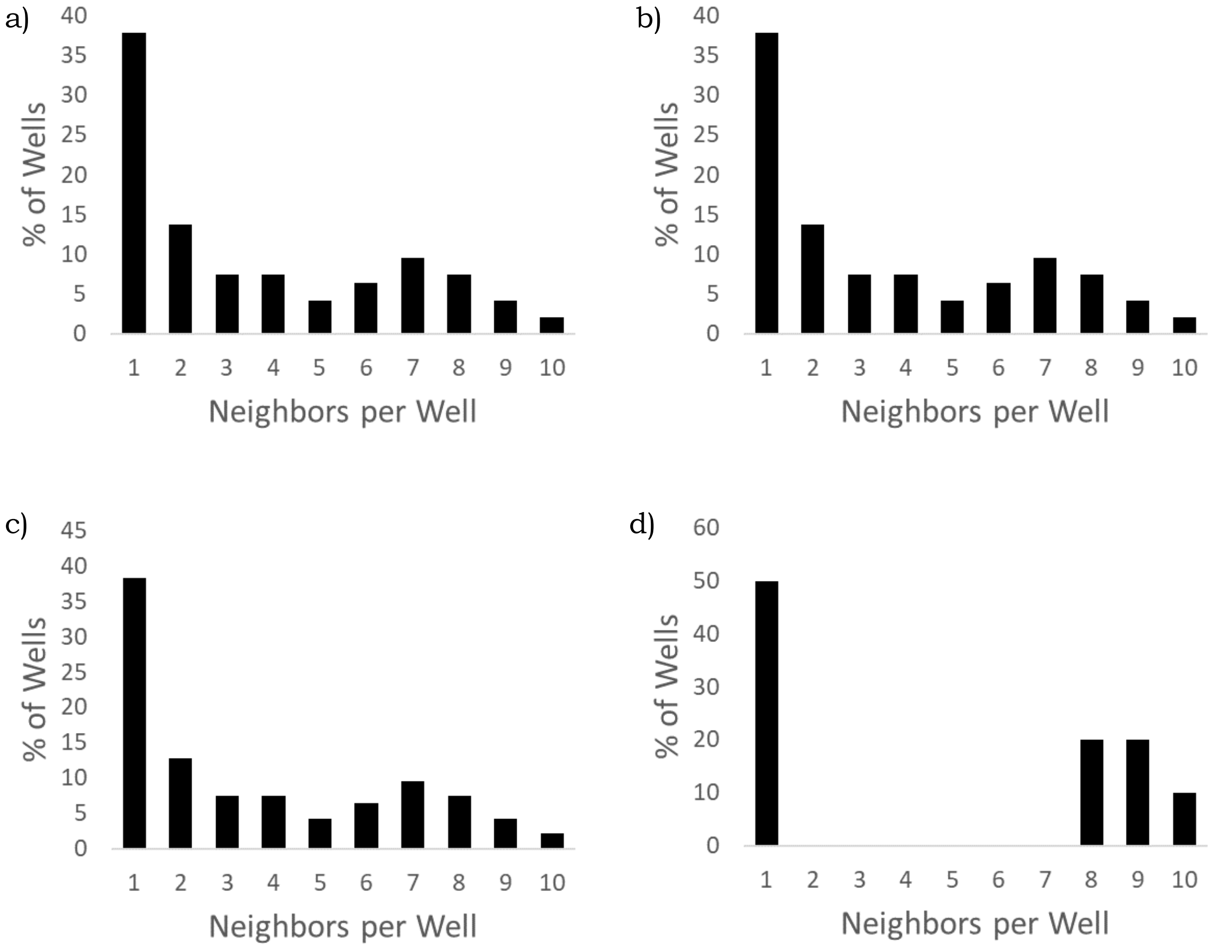


Figure S1. Neighbors per subpopulation for a) all subpopulations in heterogeneous corridor arrangement treatment b) extinct subpopulations of 90% extinction level treatment c) extinct subpopulations of 50% extinction level treatment d) extinct subpopulations of 10% extinction level treatment.

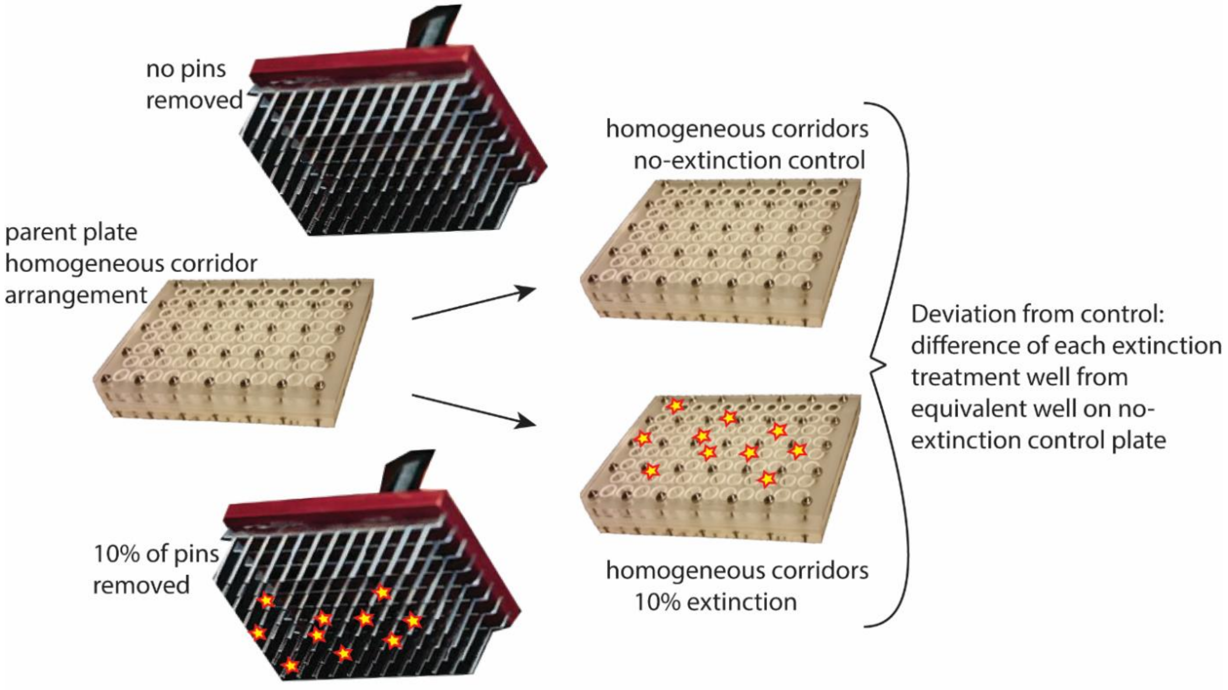


Figure S2. Response variable "Deviation from control" was calculated by subtracting fluorescence of each treatment well from the fluorescence of the corresponding well of the no extinction control. In this example, the 10% extinction treatment plate is matched to its no-extinction control plate.

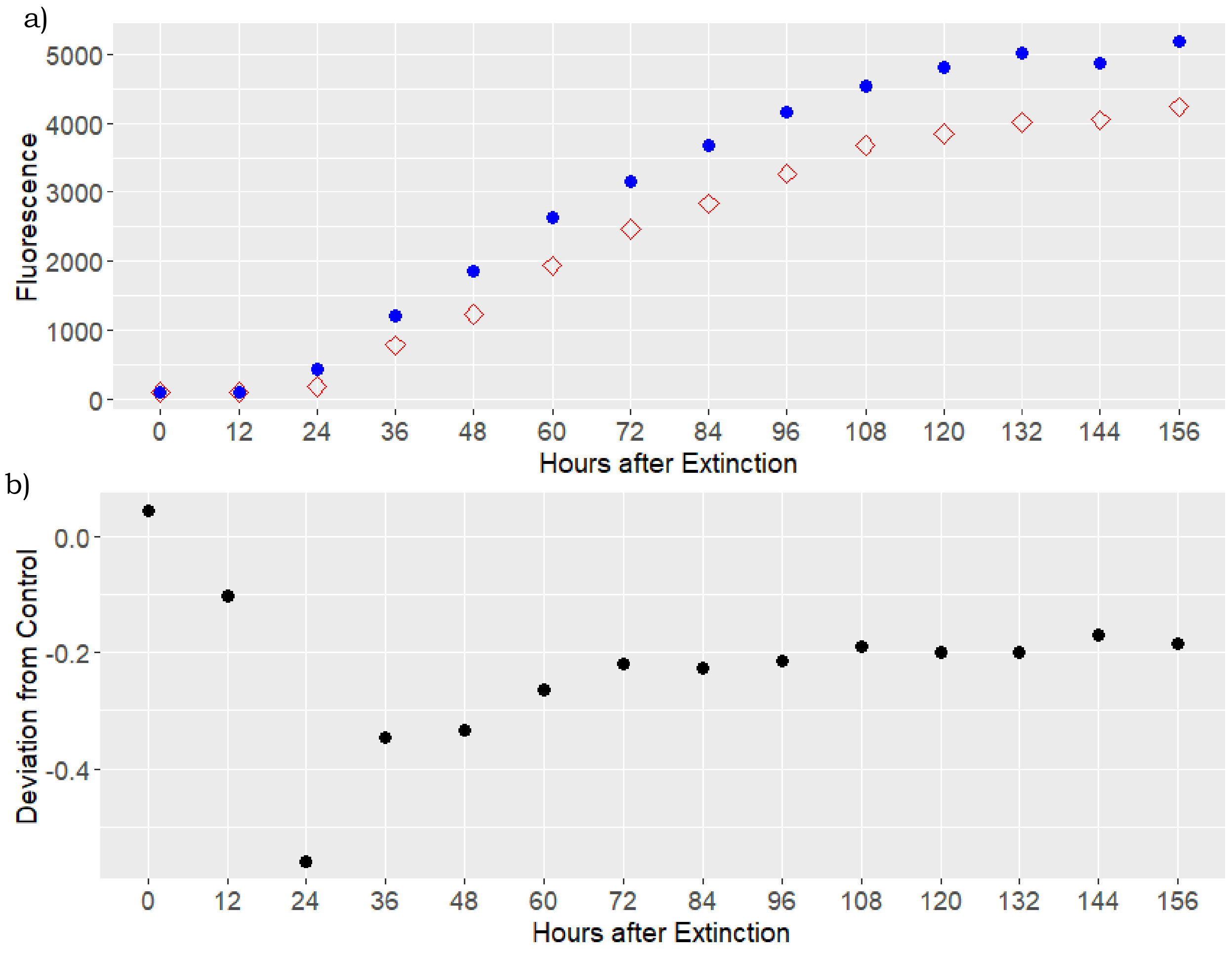


Figure S3. Example of raw data and calculation of Deviation from Control a) Time series of fluorescence in well A5 in one pair of plates with heterogeneous corridors, blue circles are well A5 in no-extinction control, red diamonds are well A5 in 10% extinction treatment b) Deviation from Control calculated by subtracting fluorescence in control well (blue circle, panel a) from fluorescence in treatment well (red diamond, panel a), divided by fluorescence in control (blue circle, panel a).
